# BEDMS: A metadata standardizer for genomic region attributes

**DOI:** 10.1101/2024.09.18.613791

**Authors:** Saanika Tambe, Oleksandr Khoroshevskyi, Sang-Hoon Park, Nathan J. LeRoy, Donald R. Campbell, Guangtao Zheng, Aidong Zhang, Nathan C. Sheffield

## Abstract

High-throughput sequencing technologies have generated vast omics data annotating genomic regions. A challenge arises in integrating this data because the associated metadata does not follow a uniform schema. This hinders data management, discovery, interoperability, and reusability. Existing tools that address metadata standardization issues are generally limited in scope and targeted toward specific data sets or types and are not generally applicable to custom schemas. To improve standardization of genomic interval metadata, we have developed BEDMS. We developed and evaluated several model architectures and trained models that achieved high performance on held-out training data. With a trained model, BEDMS provides users with predicted standardized metadata attributes that follow a standardized schema. Furthermore, BEDMS provides the ability to train custom models. To demonstrate, we trained BEDMS on three different schemas, allowing users to choose which schema to standardize into. We also deployed BEDMS on PEPhub, which provides a graphical user interface to allow users to standardize metadata without requiring any local training or software at all. In conclusion, BEDMS offers a practical one-stop solution for metadata management and standardization for genomic interval data.

## Introduction

The rapid development of high-throughput sequencing technologies has led to a surge in the volume of ‘omics data [1, 2], including genomic interval data [3]. While more data benefits research, it also leads to new challenges in data curation as research more frequently integrates data from diverse sources. A major obstacle to discovering, accessing, and integrating data is that data and associated metadata are typically formatted according to repository-specific or platform-specific schemas. Because numerous platform-specific schemas exist, ensuring that the metadata adheres to the FAIR guiding principles (Findability, Accessibility, Interoperability, Reusability) poses additional hurdles [4–6]. This challenge is particularly important for tasks that gain power from data scale, such as meta-analyses and machine learning [7].

As data generation accelerates, the challenges of metadata management will only intensify. The need to standardize metadata arises in two distinct but complementary scenarios: the use case of the data depositor, who generates and uploads the data initially to public repositories, such as NCBI GEO, and the secondary analyst, who downloads data for integrative analysis. The data depositor must grapple with standardizing metadata from its original form to fit the requirements of the target data repository. While checklists and standards encouraging FAIR principles exist for standardizing the metadata [12], depositors often prioritize other aspects over adherence to these standards. Meanwhile, the secondary analyst faces a similar challenge in merging metadata terms across diverse sources when downloading published data for re-analysis[13]. Both use cases, at opposite ends of the data cycle, share a root need: transforming non-standard metadata into standardized terms with minimal data loss.

This need is beginning to be addressed by efforts to improve discoverability and consistency of omics-related metadata [18]. For example, Cannizzaro et al. [19] used seq2seq models to map unstructured metadata into standardized metadata, and ALE (Automated Label Extraction) used heuristic and machine learning to extract standardized labels of metadata tables [17]. While these and related efforts [21] have potential to improve metadata standardization, many have been limited in number of attributes, in choice of schemas provided for standardization, in quality of software, or to specific domains [22], such as metabolomics standardizer SMetaS [23].

To address the issue for genomic region data, we have developed BEDMS – a metadata standardization tool for genomic regions. We assembled training and test datasets for a specific schema (BEDMS-condensed ENCODE schema), and then trained and evaluated 6 model architectures on that schema. We settled on a neural network architecture and used it to train separate models for this schema. We later developed training sets for two additional schemas. To provide an easy user interface to BEDMS, we deployed it in PEPhub [24], where users can use it to suggest standardized column headers as they edit BED file metadata tables. BEDMS is a step forward in helping both metadata creators and consumers by automating the process of standardizing and integrating metadata. BEDMS can be used independently through a Python package and can also be trained by users on custom schemas.

## Materials and Methods

### Overview of the approach

A sample metadata table has two types of information: attributes and values (Fig. 1A). The first step to integrating tables is to ensure that attributes match. We therefore seek a model that will take a non-standardized metadata table and provide suggestions on attributes (column headers) to comply with a standard schema. For example, for a table with columns `species and library_strategy`, standardizing under the ENCODE Project Metadata schema would suggest the attribute names `Biosample organism` and `Assay` to comply with the schema (Fig. 1B).

**Figure 1:**
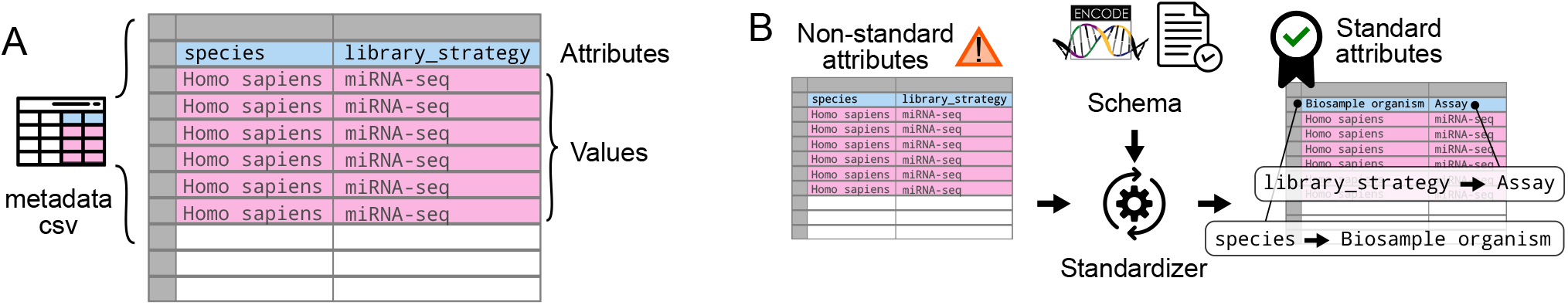
Overview of metadata standardization task and BEDMS approach. A) Sample metadata tables have two types of information. Attributes, corresponding to column headers, define the schema of a sample may define, and row values provide the actual values of those attributes. B) Schematic describing the process of suggesting new attributes that fit with a provided schema.

To achieve this, we developed six different neural network model architectures of increasing complexity. We first developed a comprehensive semi-synthetic training dataset following the ENCODE Project Metadata [25] schema, which we used to evaluate the models. After selecting the best-performing architecture for BEDMS, we built additional training datasets and trained additional models on FAIRtracks [14] and BEDbase schemas. Finally, we deployed BEDMS on PEPhub for interactive use.

### Data collection

For training, we first constructed a BEDMS-condensed ENCODE metadata schema based on the schema used for the ENCODE Project. The original metadata had 59 attributes, which we filtered to 18 attributes to develop a minimal schema. This helped adhere to FAIR principles [26] and was suitable for future automated standardization. To select these attributes, we first reduced the metadata to samples with BED (Browser Extensible Data) file type. Then, we manually refined the schema to only have attributes with biological relevance with the following steps:

1. Removing attributes such as “Audit ERROR” and “File analysis status” that were specific to the ENCODE Project and lacked general applicability.
2. Removing attributes with no values among the BED samples, such as “Read length” and “Mapped read length.”
3. Removing non-machine-learning-friendly attributes, such as “Biological Replicates,” “Technical Replicates,” where values lacked clear semantic meaning, and columns did not have exclusive values which made them unsuitable for prediction.
4. Removing attributes with constant or values, such as “File format,” “File type,” and “File format type,” which were either “bed,” “narrowPeak,” or “bed narrowPeak.”

These steps helped us generate BEDMS schemas (for ENCODE, FAIRTRACKS, and BEDbase) that would be suitable for machine learning.

We then built a comprehensive training data set through a series of manual and automated curation steps (Fig. 2A). We started from the standardized official data published from the ENCODE project (n=13,034 rows). Next, we augmented this data by of adding table rows using ontologies. For sample attributes with reasonable ontology candidates, we selected suitable ontology terms and added corresponding rows to the table (Table S1). For example, for the *Biosample organism* attribute, we added terms from the Catalogue of Life ontology [27]. Next, in the diversification step, we used ChatGPT 3.5 [28] to add synonyms, typographical errors, and related terms, based on the existing terms (See Methods). This resulted in a final table with 50,000 samples. We then generated tables from this main dataset and split these tables into training, test, and validation sets in the ratio of 8:1:1. To evaluate the model, we used three sets of test data with varying semantic similarity from the training data:

**Figure 2:**
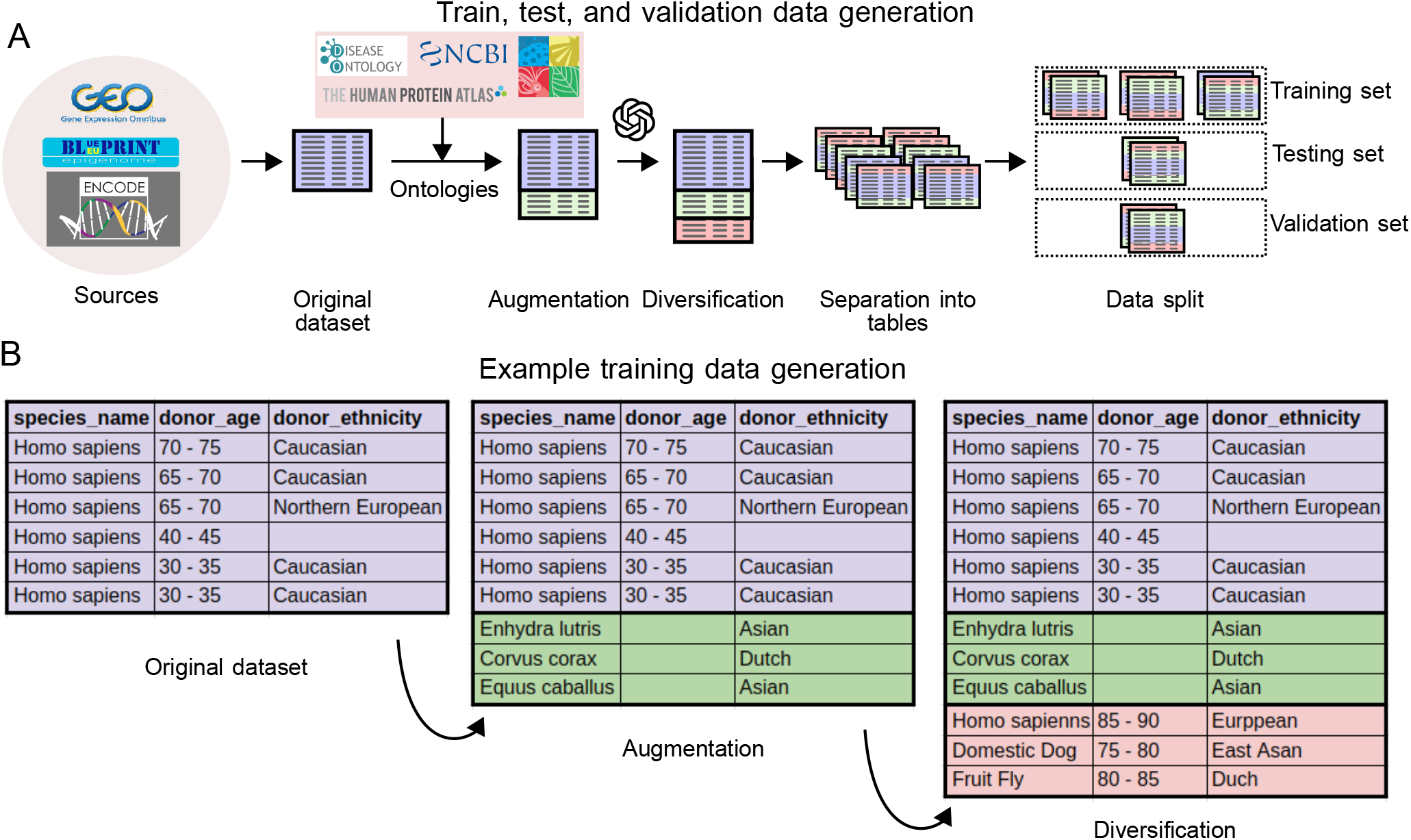
Process for constructing BEDMS training, testing and validation sets. A) Diagram showing the process of building training, testing, and validation sets. Starting from a well-annotated source, we augment with ontologies, then diversify with ChatGPT. Finally, we construct metadata tables to simulate real-world use and then split these into training, test, and validation sets. B) Concrete examples of metadata add during the training set building process.

1. Basic test set: 10% of the test data after train-test-validation split.
2. Moderate test set: Extended the test dataset with additional values sourced from the same ontologies used during training, but which were not used for training, and some values generated using ChatGPT. Approximately 40% of the values in this test were not seen by the model during training.
3. Challenging test set: Similar to the moderate test set, this set was also sourced values from the same ontologies and with values generated from ChatGPT that were not seen during training. It had 70% of the values not seen by the model during training and is meant to consist mostly of outlier or uncommon values.

These steps help the training dataset capture a more realistic set of the possible terms and errors that real world data could contain and allow it to generalize better to unseen data (Fig. 2B).

### Model architecture descriptions

We next explored 6 possible model architectures for BEDMS. We consider metadata attribute standardization as a classification task, where the input is a metadata table, and the goal is to adjust the column headers to terms from an external schema. We reasoned that two independent pieces of input information would be useful: 1) the “proposed” attribute (of the input table), and 2) the values of the attribute. We encoded the proposed attributes as either one-hot encoded vectors or Sentence Transformer embeddings [29], and values as either one-hot encoded vectors, bag of words vectors, or Sentence Transformer embeddings (Fig. 3A). For all Sentence Transformer embeddings, we used model all-MiniLM-v2-L16. We developed six neural networks (NNs) architectures that use different encoding types for each of the inputs (Fig. 3B). The overall architecture of these six models is similar, but they differ in the structure and types of input encoding, with increasing complexity:

**Figure 3:**
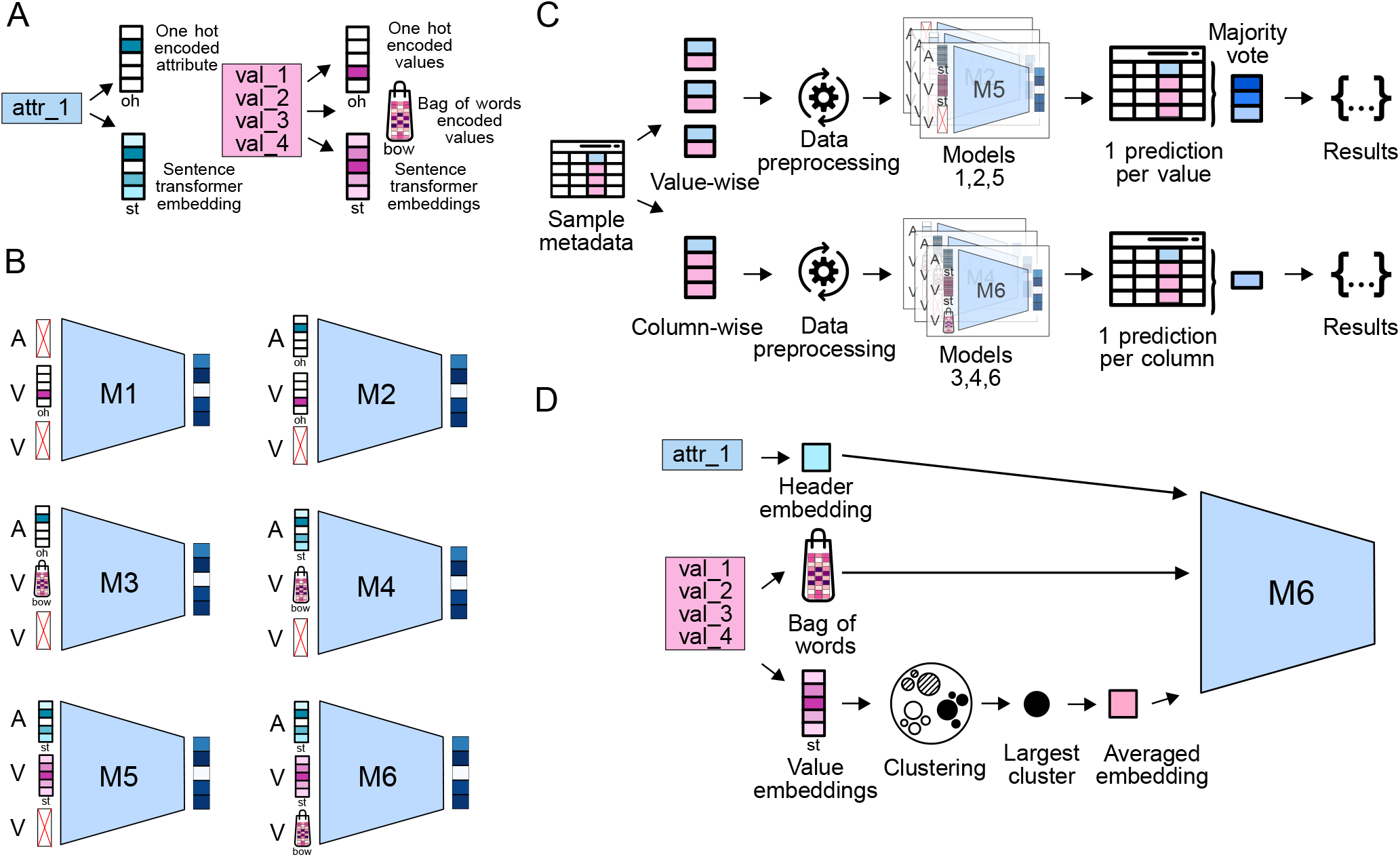
Candidate BEDMS model architectures, input encoding methods, and training procedures. A) Schematic showing different ways to represent the attributes and values of the input metadata. B) Diagram showing which input encodings are used for each of 6 candidate models. C) Data preprocessing for Model 6 showing how value embeddings are pooled. D) Overall model processing procedure, depicting the differences between value-wise models (1,2 and 5) and column-wise models (3, 4, and 6).

Model 1: Model 1 provides a baseline for comparing the other models. It considers only the values in the table, ignoring the suggested attribute name from the input. Each value in the training metadata is one-hot encoded.

Model 2: Model 2 builds on Model 1 by adding the attribute as a one-hot encoded vector. The values and attributes are separately one-hot encoded and fed into the model as two distinct input layers which are then concatenated before the combined output layer. This architecture allows the neural network to learn associations between values and their corresponding input attributes.

Model 3: Model 3 replaces the one-hot value vector into a Bag of Words (BoW) encoding. Like Model 2, these get concatenated before the combined output layer. Unlike models 1 and 2, which operate the level of individual values, Model 3 uses an aggregate representation of values, so it is trained on the entire attribute column. Because the individual values are no longer independent, this allows the model to consider the frequency of the terms simultaneously, instead of treating each term as a distinct entity.

Model 4: Model 4 builds on Model 3 by changing the attributes embeddings from a one-hot vector into a Sentence Transformer embedding. This allows the model to learn from the semantic meaning of the input attributes [30].

Model 5: Model 5 builds on Model 4 by embedding not just the attribute, but also the values with the Sentence Transformer. We realized that the semantic meaning of the values may also be useful. Because Model 5 uses individual sentence transformer embeddings for each value, it is a value-level model like models 1 and 2, rather than a table-level model.

Model 6: Finally, Model 6 seeks to combine the advantages of both BoW encoding and sentence transformer embeddings. It combines the approaches used in Models 4 and 5 to come up with our most complex model. The encoding approach for Model 6 is also different from the other models. It uses the Sentence Transformer attribute embedding from Models 4 and 5, and the Sentence Transformer value embedding from Model 5, but adds in the Bag of Words value vector from Model 4. This combination of encodings provides the model with both a semantic representation, and a vocabulary frequency representation.

### Data preprocessing

The 6 models follow similar general preprocessing steps to maintain uniformity where possible, but vary where necessary to accommodate the encoding differences (Fig. 3C). The most important difference is in data input: Models 1,2, and 5 encode individual values, and thus provide value-wise predictions. To convert the value-wise predictions into an overall prediction for an entire table, for these models, we simply use a majority vote of the value-wise predictions. In contrast, models 3,4, and 6 encode an entire sample table into a single prediction, thus providing column-wise suggestion out-of-the-box. Another important difference is that in Model 6, we process the sentence transformer attribute embeddings differently. Because this is a column-level model, we require a way to pool the embeddings across values (Fig. 3D). To pool the embeddings, we first cluster them. Then we take the largest cluster and compute its average. That is used as the embedding for the value. For each column, there is a single sentence transformer embedding generated for the header, there is one embedding generated for the values and one bag of words generated for the values. For all models, after encoding, we converted the data into PyTorch tensors which were used as inputs to the model.

### Training procedure

We first performed hyperparameter tuning using Optuna [31]. We chose Cross Entropy Loss function and Adam optimizer for all our models. We used batch training to optimize memory usage and trained the models over a maximum of 20 epochs and implemented early stopping.

### Model evaluations and comparison

The models were evaluated using overall and class-specific accuracy, precision-recall, and Receiver Operating Characteristic (ROC) curves.

### Additional models

After selecting the best-performing architecture, we then built similar training data sets and training models for two additional schemas, one for the FAIRtracks schema, and one for the BEDbase schema.

## Results

### Test accuracy comparison and evaluation

We evaluated the models on 3 test sets of increasing difficulty: easy (no unseen values), moderate (40% unseen values) and challenging (70% unseen values). The easy and moderate tests are akin to what we expect the models to encounter in real-world scenarios, where we expect many of the values to match something in the training set. The challenging test data was designed to assess the models’ robustness on highly unexpected data and outliers. As expected, the models performed best on the easy test set, followed by the moderate and then challenging tests in terms of overall accuracy (Fig. 4A) and F1 score (Fig. 4B). We also observed a general trend where the more complex models (with additional input variables and more sophisticated encodings) outperformed the simpler models. On the easy test, Model 6 achieved the highest accuracies, whereas on the moderate and challenging tests, Model 5 performed the best. Overall, Models 5 and 6 both had substantially superior performance to the other models. This is likely due to their use of sentence transformers for both headers and values, allowing these models capture semantic meaning of both values and proposed attributes.

**Figure 4:**
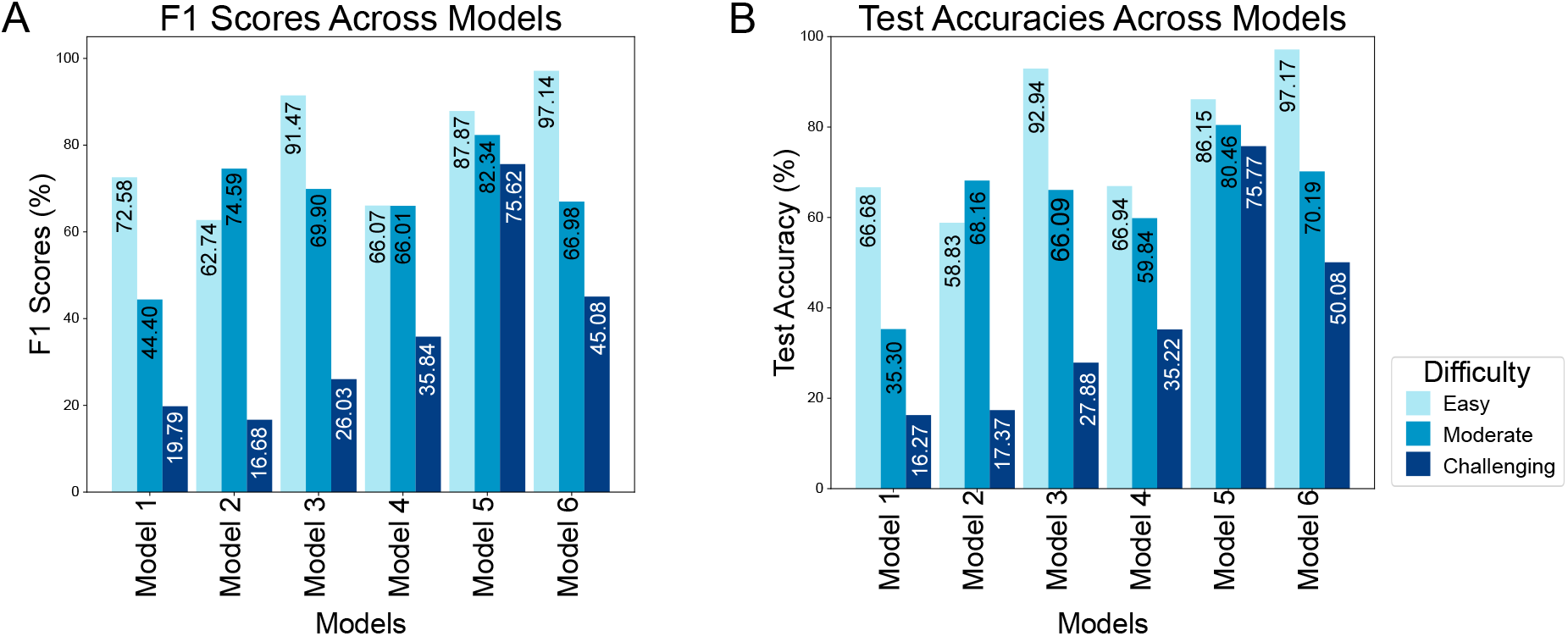
Results of evaluation tests on ENCODE-trained BEDMS model. A) Bar graph showing the F1 scores across all model architectures for three different test sets - easy, moderate, challenging. B) Bar graph showing the test accuracies across all model architectures for three different test sets - easy, moderate, challenging.

### Class-specific accuracy assessment

In our assessment of class-specific accuracies, we noticed consistency among models, where they tend to perform well for some classes as opposed to the others. This implies difficulty differences among classes, which could be caused by the values for some attributes being more semantically meaningful than others. To investigate this further, we visualized model accuracy by class (Fig. 5A). We produced confusion matrices (Fig. S1) and columnwise ROC curves (Fig. S2) for all combinations of model and test set. We then further explored how the sentence transformer embeddings of the values for each are embedded using t-SNE (Figure 5B). Values in easy-to-predict columns, such as *Biosample organism*, the value embeddings clustered tightly, whereas, harder-to-redict attributes such as *Assay* do not (Figure 5B). When unseen values for diffuse attributes are provided, this could explain why the model underperforms. To confirm that our clustering procedure used in Model 6 is not leading to strange embeddings that could impact model performance, we also visualized embeddings of the averaged values following the Model 6 pooling procedure, which confirmed that these embeddings have similar characteristics to the individual value embeddings, providing confidence that this procedure is robust (Figure 5C).

**Figure 5:**
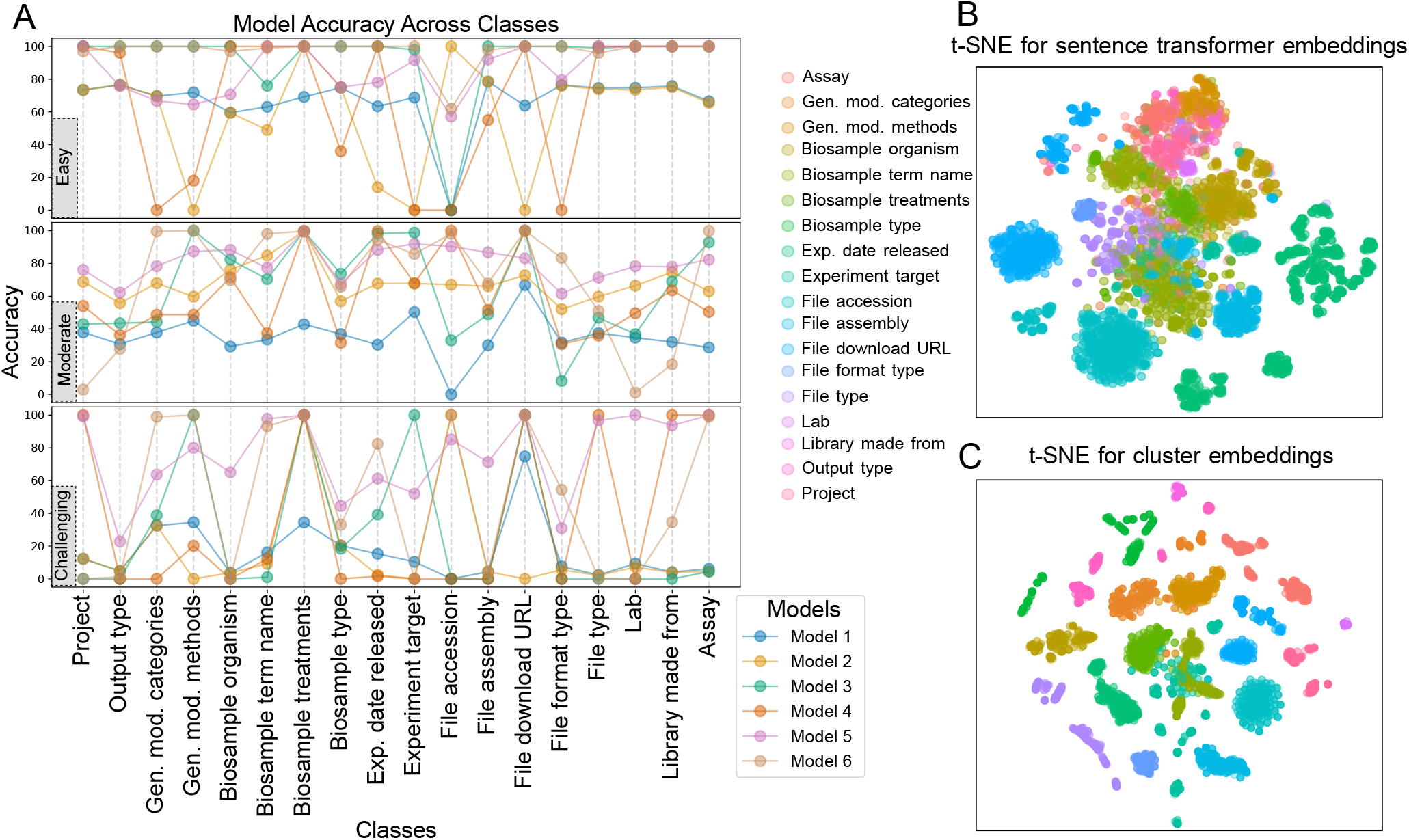
Columnwise results and embeddings. A) Dotplot showing columnwise accuracy of the 6 candidate model architectures for 3 evaluation sets - easy, moderate, and challenging. B) t-SNE plot showing the clustering of value embeddings for the BEDMS-condensed ENCODE metadata schema. C) t-SNE plot showing the clustering of value embeddings for the BEDMS-condensed ENCODE metadata schema after preprocessing according to the model 6 architecture.

Based on our findings, we selected Model 6 as the final BEDMS architecture, as it provides the best performance on the most realistic datasets and good performance on outlier results and, since it operates at the column level, also provides potential for performance benefits over the similarly-performing Model 5. The final model architecture for BEDMS thus consists of 3 independent inputs, with the value represented as both a Bag of Words and Sentence Transformer embedding, and the attribute as a Sentence Transformer Embedding (Fig. 6A). These input layers are fully connected independently to a first hidden layer, which is concatenated and then run through two more fully connected hidden layers. Model 6 performance in detail shows almost perfect performance on the easy test set, as well as good overall performance on the moderate and challenging test sets, with specific problematic attributes that can probably be improved with future tuning (Fig. 6B).

**Figure 6:**
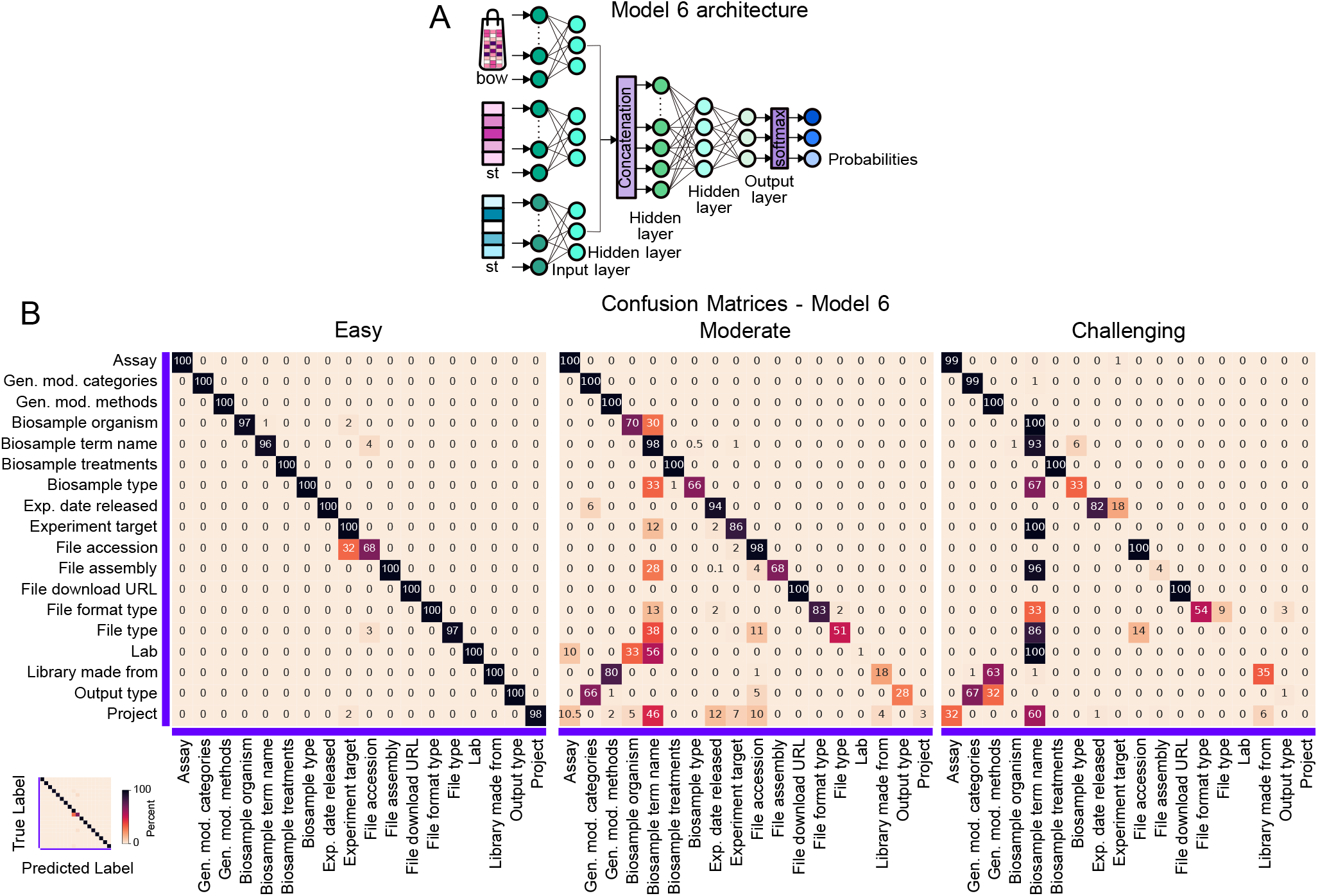
Model 6 test results. A) Detailed model 6 architecture. Model 6 takes three inputs (bag of words encoded values, sentence transformer embedded values, sentence transformer embedded headers), concatenates them, and provides the output. B) Confusion matrices of model 6 architecture for three different tests-easy, moderate, challenging.

### PEPhub deployment and user experience workflow

To make BEDMS usable by others, we deployed it in two ways. First, it can be downloaded as a Python package. You can use the 3 pre-trained models we described for the 3 schemas, which are available on Hugging Face to standardize your attributes. Furthermore, we also deployed the BEDMS metadata standardizer on PEPhub for users to standardize their metadata attributes using a web interface. The PEPhub database allows a user to store and manage their biological metadata in the form of a Portable Encapsulated Project (PEP). A basic PEP consists of tabular metadata stored in csv and JSON format. To standardize metadata on PEPhub, a user selects the sample metadata table, selects a schema from the available options, and then BEDMS suggests changes to each attribute in the user’s sample metadata, along with a confidence level indicating how confident BEDMS is in the accuracy of the suggestion. The user then can approve and apply the changes to their sample metadata or reject them (Fig. 7). On approval, the attributes are standardized as per the suggestions. On rejection, no changes are made to the sample metadata and the user can choose to try out a different schema.

**Figure 7:**
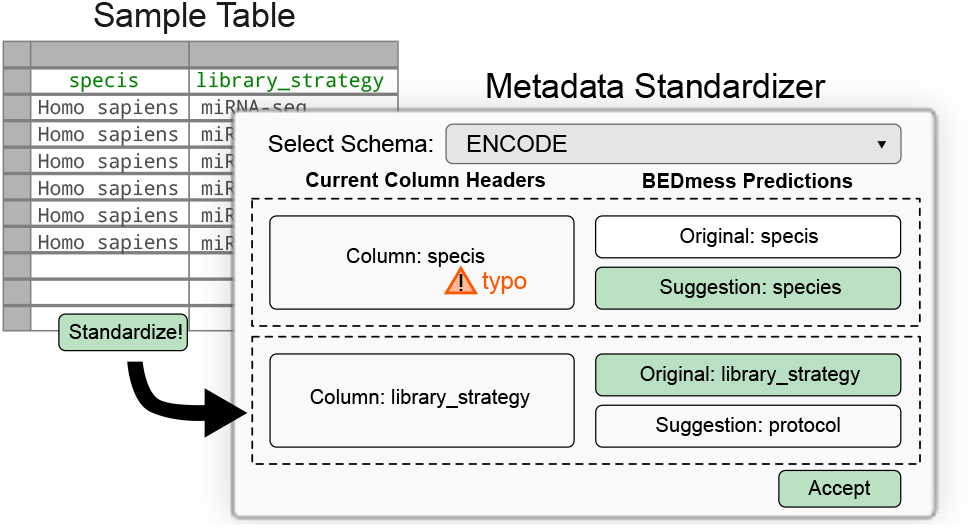
Deployment of BEDMS on PEPhub. Diagram showing the user experience of BEDMS on PEPhub. The user provides a sample table to BEDMS and receives suggestions for each attribute. If they have made a spelling error, BEDMS will suggest an attribute name with the correct spelling or if the attribute name is not matching the schema of user’s choice, BEDMS will provide a suggestion that fits with the schema. They can select from the options or choose to keep the original one.

## Discussion

Metadata standardization is crucial to the FAIR guidelines for data sharing, but there are still too few tools that help researchers standardize metadata from diverse sources.

This is partly due to the difficulty and diversity of the challenge, which requires tailor-made solutions for different data types. Existing efforts in this direction are typically restricted to particular data types [23], and there are few efforts specifically designed for genomic/epigenomic metadata.

To address this issue for this data type, BEDMS provides a model architecture and training approach to train custom models for standardizing metadata of BED files. BEDMS can be used to make non-standardized data fit a schema, or it can also be used to translate standardized data from one schema to another. In this paper, we have described various approaches we pursued to develop BEDMS and its applications. Key contributions include: First, we developed 6 different neural network models of increasing complexity and systematically evaluated them to ascertain which models perform the best for our specific task. Second, we proposed a novel approach to solving the metadata standardization problem by combining sentence transformers and bag of words encoding. This method captures both the semantic context and the word frequency in the metadata, improving robustness and accuracy. Third, we deployed our chosen model into the PEPhub web interface to make it more accessible to users in a graphical interface. Finally, our Python module provides tools for users to train custom models for custom schema.

Overall, BEDMS establishes a framework for using machine learning for automated metadata attribute standardization for genomic region set data. Future effort in this direction can alleviate some of the limitations of our current approach. BEDMS currently only standardizes metadata attributes; we did not address the related issues of standardizing values themselves or project-level annotation terms or tags. BEDMS is also limited to BED file metadata, and it requires a separate model for each schema. With more research it may be possible to train a single foundational model capable of standardizing metadata across schemas and data types.

In conclusion, BEDMS presents a robust framework for automating metadata standardization using advanced neural network models. BEDMS has significant potential in facilitating data management for genomic and epigenomic metadata.

## Supplemental material

### Using ChatGPT for training set diversification

We used ChatGPT to diversify the values generated for our training data sets. For this, we provided examples of the acceptable values that would be suitable for the attributes and prompted ChatGPT to generate synonyms or similar alternatives. We noticed that ChatGPT provided high quality suggestions only for approximately the first 100 values that it would generate following which the values would become highly general or reptitive. Hence, we specified the limit for each query ‘s results to be 100 values to maintain the quality of data. Here are some example prompts that we used:

For generating dates in different formats: “generate 100 dates in format : 15 Mar. 1990,” “generate 100 dates in different formats.”

For generating experiment targets: “provide examples of targets like: RXRA-human, H3K9me3-human, XRN2-
human APEH-human”

For generating spelling errors: “can you provide 100 spelling errors for these genetic modification categories: interference, deletion, insertion, interference”

For generating spelling errors in organism names: “Can you generate 100 spelling mistakes of scientific names of organisms”

We used these kinds of prompts across all attributes for all three schemas that BEDMS standardizes into.

**Figure S1:**
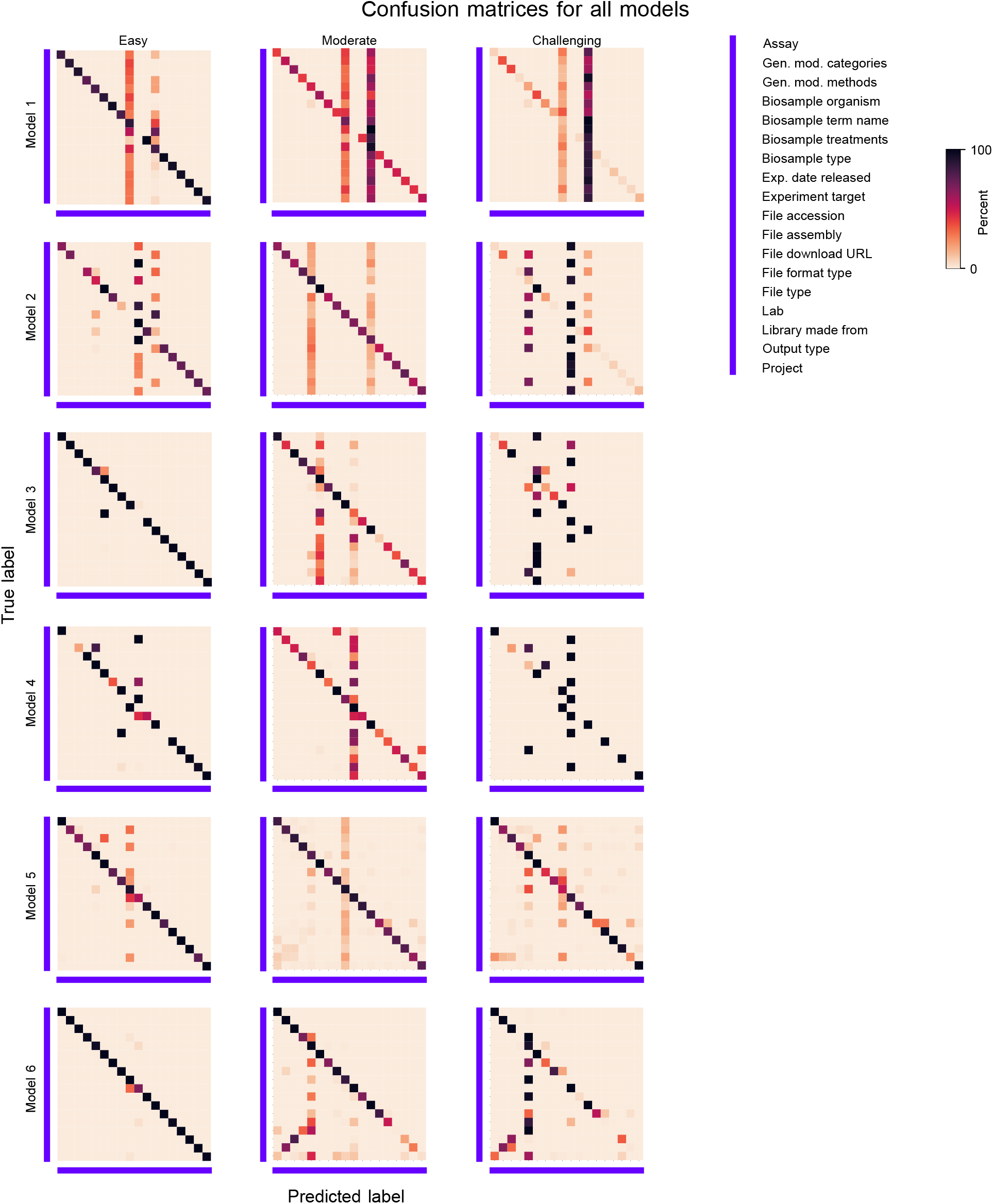
ROC curves for column-wise accuracy evaluation. ROC curves for each of the attributes in the ENCODE schema, showing how the 6 candidate BEDMS architectures perform on each column for each of the 3 test sets, easy, moderate, and challenging..

**Figure S2:**
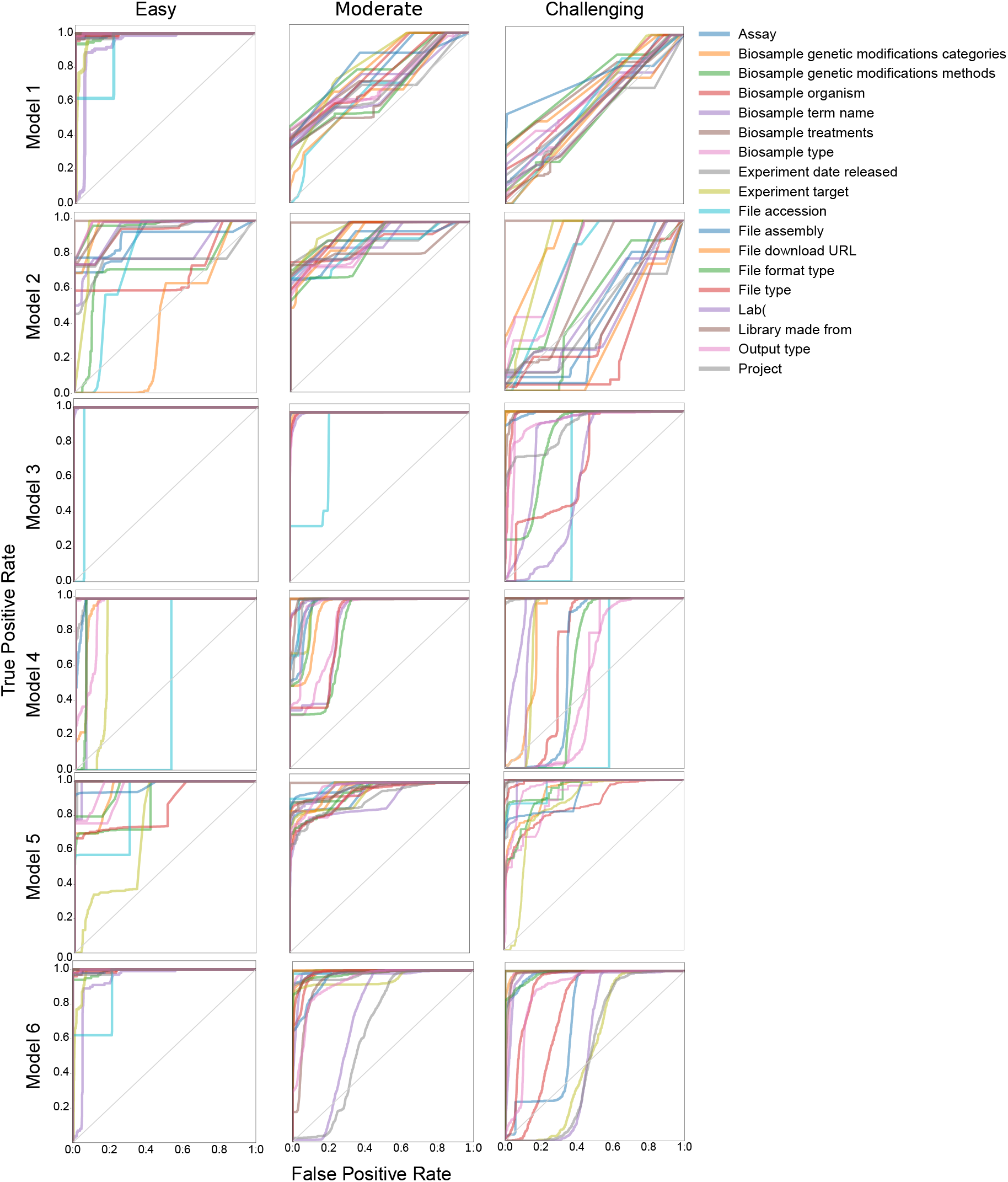
ROC curves for column-wise accuracy evaluation. ROC curves for each of the attributes in the ENCODE schema, showing how the 6 candidate BEDMS architectures perform on each column.

**Table S1:** List of ontologies. Table listing attributes for the schemas where we used external ontologies.

| Schema | Attribute | Source | Reference |
| --- | --- | --- | --- |
| ENCODE | File assembly | NIH NLM Genome Reference | OLeary 2024 |
| ENCODE | Biosample term name | Human Protein Atlas | Uhlen 2015 |
| ENCODE | Biosample term name | Publication | Hatton 2023 |
| ENCODE | Biosample organism | Catalogue of Life | Banki 2024 |
| FAIRTRACKS | species_id | NCBI Taxonomy | Schoch 2020 |
| FAIRTRACKS | species_name | Catalogue of Life | Banki 2024 |
| FAIRTRACKS | donor_health_status | Disease Ontology | Bernasconi 2019 |
| FAIRTRACKS | donor_health_status | Alliance of Genome Resources | Alliance 2024 |
| FAIRTRACKS | sample_type_term_label | Human Protein Atlas | Uhlen 2015 |
| FAIRTRACKS | sample_type_term_label | Publication | Hatton 2023 |
| FAIRTRACKS | phenotype_term_label | Disease Ontology | Bernasconi 2019 |
| FAIRTRACKS | phenotype_term_label | Alliance of Genome Resources | Alliance 2024 |
| BEDBASE | genome | NIH NLM Genome Reference | OLeary 2024 |
| BEDBASE | species_name | Catalogue of Life | Banki 2024 |
| BEDBASE | species_id | NCBI Taxonomy | Schoch 2020 |
| BEDBASE | phenotype | Disease Ontology | Bernasconi 2019 |
| BEDBASE | phenotype | Alliance of Genome Resources | Alliance 2024 |
| BEDBASE | cell_type | Publication | Hatton 2023 |
| BEDBASE | cell_line | Human Protein Atlas | Uhlen 2015 |
| BEDBASE | tissue | Publication | Hatton 2023 |
| BEDBASE | antibody | NCBI Entrez | Entrez 2010 |

